# Imaging mass spectrometry and shotgun proteomics reveal dysregulated pathways in hormone induced male infertility

**DOI:** 10.1101/2020.02.03.931931

**Authors:** Shibojyoti Lahiri, Lena Walenta, Wasim Aftab, Leena Strauss, Matti Poutanen, Artur Mayerhofer, Axel Imhof

## Abstract

Spermatogenesis is a complex multi-step process involving intricate interactions between different cell types in the male testis. Disruption of these interactions results in infertility. Combination of shotgun tissue proteomics with MALDI imaging mass spectrometry is markedly potent in revealing topological maps of molecular processes within tissues. Here, we use a combinatorial approach on a characterized mouse model of hormone induced male infertility to uncover misregulated pathways. Comparative testicular proteome of wildtype and mice overexpressing human P450 aromatase (AROM+) with pathologically increased estrogen levels unravels gross dysregulation of spermatogenesis and emergence of proinflammatory pathways in AROM+ testis. *In situ* MS allowed us to localize misregulated proteins/peptides to defined regions within the testis. Results suggest that infertility is associated with substantial loss of proteomic heterogeneity, which define distinct stages of seminiferous tubuli in healthy animals. Importantly, considerable loss of mitochondrial factors, proteins associated with late stages of spermatogenesis and steroidogenic factors characterise AROM+ mice. Thus, the novel proteomic approach pinpoints in unprecedented ways the disruption of normal processes in testis and provides a signature for male infertility.

---

Infertility is a major health condition that affects around 15% of couples of reproductive age worldwide. About a third of these occurrences are due to male infertility. During the recent decades male reproductive health has declined in many parts of the world contributing, cumulatively, to about 65% of all infertility cases worldwide(Li, Strauss et al., 2006, Singh & Singh, 2017). A variety of chromosomal anomalies like Klinefelter syndrome(Gravholt, Chang et al., 2018), Y-chromosome deletions(Song, Chiba et al., 2016) and mutations in spermatogenic genes(Bracke, Peeters et al., 2018) have been attributed to reduced fertility in men. Apart from these, a number of anatomical and physiological features have been associated with infertility in men, but precise causative molecular mechanisms remained elusive so far. This has resulted in a substantial percentage of male infertility cases to be classified as idiopathic (Singh & Singh, 2017) and hence, subject to only empirical treatments. Systematic understanding of spermatogenesis at the molecular level is therefore imperative in attempts to tackle the aforesaid condition.

Spermatogenesis is a process that results in the production of fully differentiated male gametes. It involves a series of intricate interactions between different cell types and molecules within the testis. A number of genetic, epigenetic and environmental factors have been shown to affect spermatogenesis leading to reproductive failure in men (Neto, Bach et al., 2016). In majority of cases, the testis is the principal organ giving rise to abnormalities leading to male infertility. High levels of intratesticular testosterone are required for testis to produce spermatozoa and impaired sperm production can result from increased circulating estrogen levels (Ciaccio LA, 1978, Jones, Fang et al., 1978, Kalla, Nisula et al., 1980). To simulate such a hormonal imbalance a transgenic mouse model overexpressing the human P450 aromatase (AROM+) gene has been established (Li, Nokkala et al., 2001, Li et al., 2006). These mice have an increased E2/T ratio and develop a progressive male infertility phenotype. The disease model has so far been only characterized on a histological and phenotypic level resulting in the identification of a limited number of affected pathways (Li et al., 2001, Li et al., 2006, Strauss, Kallio et al., 2009). Detailed molecular description of the system is still lacking, which results in a limited understanding of pathways involved. Large scale -omics techniques allow a much deeper molecular understanding of the pathways and proteins involved in the phenotype. So far, most proteomic studies have focussed on sperm or seminal fluids (Panner Selvam & Agarwal, 2018). Only very few proteomic studies of the testis have been performed, which were in addition severely limited in terms of the number of proteins identified. This, along with the lack of spatio-temporal information have led to very inadequate understanding of the functional network of these proteins in spermatogenesis (Liu, Hu et al., 2013, MacLeod & Varmuza, 2013). Present day shot gun proteomic studies of complex organs can identify a large number of proteins. However, they inherently bear the disadvantage of loss of spatial information of the identified proteins. To overcome this inherent problem for a better understanding of spermatogenesis and male infertility, we first performed a deep shot gun proteomic study of testis from 11 month old AROM+ mice and compared them to wild type controls. The information about the identified peptides gathered this way was then used to learn about their *in situ* distribution using MALDI-IMS. The aforesaid exhaustive studies in combination with a newly developed data analysis tool described here provide an in-depth understanding of molecular processes affected in the testis during hormone induced and inflammation associated male infertility.

## Results

### Characterization of cellular pathways and proteins upon alteration of the E2/T ratio in testis

Mice overexpressing aromatase (AROM+) are known to exhibit symptoms of human male infertility(Li et al., 2006). The mice show cryptorchidism, Leydig cell hyperplasia and a general lack of reproductive fitness. As the processes dysregulated in these animals are not entirely clear, we performed an extensive proteomic analysis of testis tissues from WT and AROM+ mice (Figure 1A) at the age of 11 months.

**Figure 1:**
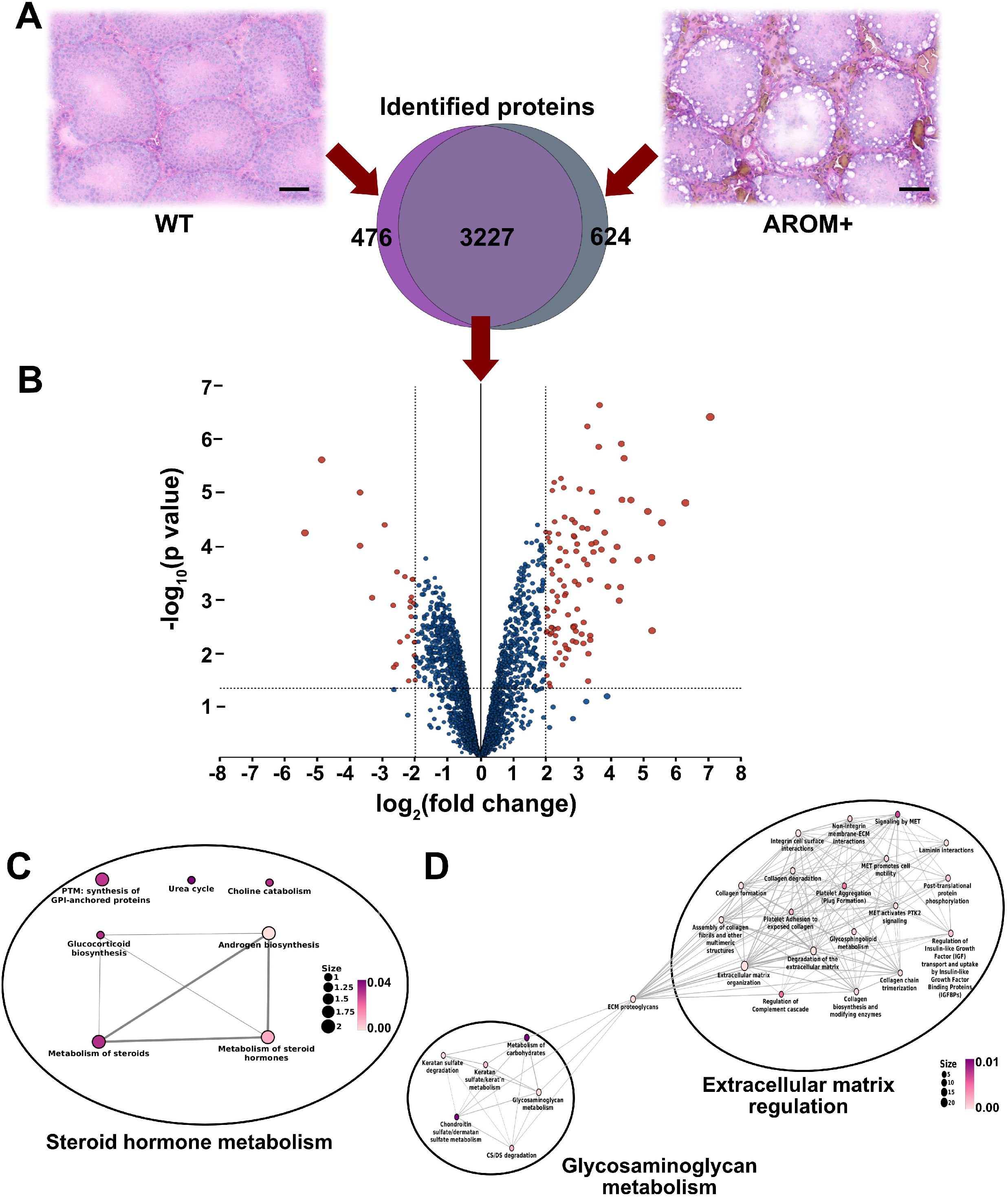
Proteomic analysis of WT and AROM+ testes. **(A)** The upper panel shows HE stained tissue sections of WT and AROM+ testis. Scale bar – 100 μm. Venn diagram in the lower panel shows that 3227 proteins were identified in both WT and AROM+. Additionally, 476 and 624 proteins were detected exclusively in WT and AROM+ respectively. **(B)** Volcano plot generated from relative abundance of the shared 3227 proteins after statistical analysis by LIMMA moderated t-test. The filled brown circles represent the significantly enriched proteins in both WT (negative fold change) and AROM+ (positive fold change). **(C)** Statistically significant molecular pathways generated by enrichment analysis using hypergeometric distribution model represented by the WT enriched proteins (Supplementary file 2). **(D)** Statistically significant molecular pathways using enrichment analysis (pathways having the lowest 25 p-values generated using hypergeometric distribution model are displayed here) represented by the AROM+ enriched proteins (Supplementary file 3) (Data from 3 mice for each group WT and AROM+ was considered for the statistical analyses) The colour code is representative of the statistical significance and is displayed according to the adjusted p-values generated using hypergeometric distribution model (as indicated by the magenta-white gradient bar). The size of the nodes in the network (scale represented by numbered black solid circles) are representative of the number of genes involved in a particular pathway within the network.

A total of 4327 proteins were identified, among which, 476 and 624 proteins were exclusively detected in the WT and AROM+ respectively (Fig 1A Venn diagram, supplementary file 1). A subsequent statistical analysis of the shared 3227 proteins revealed a further significant enrichment (> 4 fold in at least 2 out of 3 animals) of 27 proteins in the healthy mice as compared to 75 proteins in the AROM+ system (Figure 1B, brown filled circles, supplementary file 1). To identify the dysregulated cellular processes upon overexpression of aromatase, we analysed the molecular pathways to which these proteins contribute. Results reveal that regulation of androgen biosynthesis and metabolism of steroid hormones are the most highly enriched processes in WT mice (Figure 1C, supplementary file 2). Proteins involved in these pathways such as Cyp17a1 and Hsd17b3 (Supplementary file 2) have previously been shown to be severely reduced in AROM+ testis (Strauss et al., 2009). Apart from proteins involved in the above mentioned pathways, a number of other proteins were significantly enriched in WT mice. Presence of proteins like the testis isoform of Estrogen sulfotransferase, sperm mitochondrial-associated cysteine-rich protein, axonemal dynein heavy chain 12 and 17, phosphoglycerate kinase 2 and tubulin polyglutamylase TTLL5, testis specific cytochrome C, male-enhanced antigen 1, among others (Supplementary file 1), suggest dysregulation of several late stage spermatogenesis related phenomena in AROM+.

Activation of numerous biological processes are observed in the testes of AROM+ animals. Emergence of extracellular matrix (ECM) organization and disassembly among the significantly enriched processes suggest an induction of pro-inflammatory pathways upon alteration of E2/T ratio (Figure 1D, supplementary file 3). Occurrence of regulation of the complement cascade (Figure 1D), neutrophil degranulation and platelet aggregation among the enriched processes in AROM+ also reflects the induction of a substantial immune response. Pathway analysis shows that arrangement of extracellular organisation is mediated mainly via regulatory pathways of collagen, the main protein component of the ECM (Fig 1D, black oval labelled Extracellular matrix regulation, supplementary file 3). In addition, metabolism of glycosaminoglycan and its constituents, other prominent components of ECM, are found to be enriched in AROM+ (Fig 1D, black oval labelled Glycosaminoglycan metabolism, Supplementary file 3). Formation of a distinct cluster involving degradation of different glycosaminoglycans and its associated metabolic processes suggests their prominent contribution in development of the AROM+ phenotype.

### Molecular pathways severely affected upon increased E2/T ratio

In this report, proteins detected exclusively in either WT or AROM+ testis provide valuable insights into the most predominant biological processes that are dysregulated by an elevated E2/T ratio. Pathway analyses show that proteins exclusively detected in WT form 3 distinct clusters (Figures 2A, B and C). The first cluster represents various pathways related to transcription initiation and elongation (Figure 2A, Supplementary file 4), whereas the second cluster reveals processes involved in transcription termination and RNA maturation (Figure 2B, Supplementary file 4). This is in agreement with the substantial loss of germ cell differentiation and proliferation, known to be transcriptionally highly active. The third cluster (Figure 2C, Supplementary file 4) reveals mitochondrial processes to be predominant in the WT. Absence of these pathways in AROM+ along with its fertility defects suggest that higher estrogen levels impair mitochondrial function, potentially being largely responsible for the impaired steroidogenesis in the AROM+ testis.

**Figure 2:**
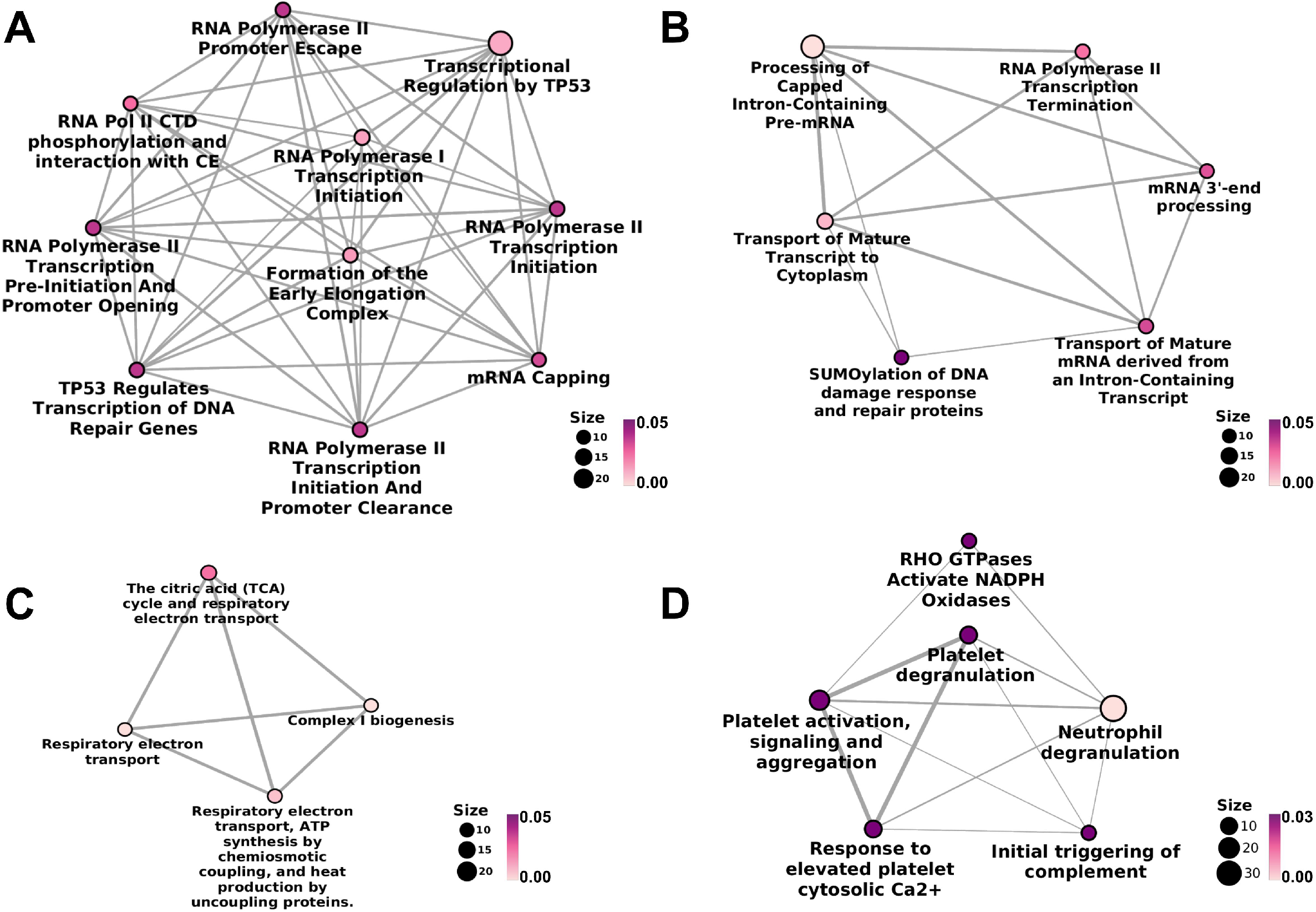
Pathways exclusive to WT and AROM+ testes. **(A)** Significant molecular pathways associated with early phases of transcription, detected exclusively in WT. **(B)** Pathways associated with later stages of transcription that are detected only in the testes of healthy animals. **(C)** Mitochondrial pathways observed solely in WT. **(D)** Inflammatory pathways observed only in 11 month old AROM+ mice, which are otherwise not observed in healthy conditions (n = 3 mice for each group WT and AROM+) The colour code is representative of the statistical significance and is displayed according to the adjusted p-values generated using hypergeometric distribution model (as indicated by the magenta-white gradient bar). The size of the nodes (scale represented by numbered black solid circles) are representative of the number of genes involved in a particular pathway within the network.

The most significant molecular pathways that emerged by analysing proteins identified exclusively in AROM+ are related to immune response (Figure 2D, Supplementary file 5). Platelet activation, degranulation and other platelet dependant regulatory processes along with neutrophil degranulation suggest the triggering of severe inflammatory and immunomodulatory activities in the testis of 11 month old AROM+ mice. Higher levels of proteins associated with ER and golgi functioning is suggestive of cellular stress in AROM+ (Supplementary file 5).

### In situ proteomic map parallels the morphological change in AROM+ testis

Increase in estrogen levels has been found to associate with cell type specific effects in male testis. Prominent fibrosis has also been observed in the interstitial spaces and tubular walls of the testis in aged AROM+ mice (Li et al., 2006). To generate a molecular map that reflects the morphological changes in AROM+ testis and to get a better understanding of the distribution of particular proteins and peptides within the tissue, we generated peptide maps of the testis from MALDI-IMS measurements by *in situ* digesting WT and AROM+ mice testis sections with trypsin. An unbiased hierarchical clustering of the acquired IMS data enabled us to distinguish different regions of the testis solely based on spectral features.

Results show two distinct types of primary clusters in the segmentation map of both WT and AROM+ represented by the brown and dark blue patches in figures 3A and 3B. The brown cluster represents peptides distinct to certain testicular seminiferous tubules (Fig 3A), which are greatly reduced in AROM+ (Fig 3B) derived tissue (~ 52% in WT as compared to 9.5% in AROM+). This peptide signature is in line with the observed overall disruption of tubular morphology in AROM+ testis(Li et al., 2006).

**Figure 3:**
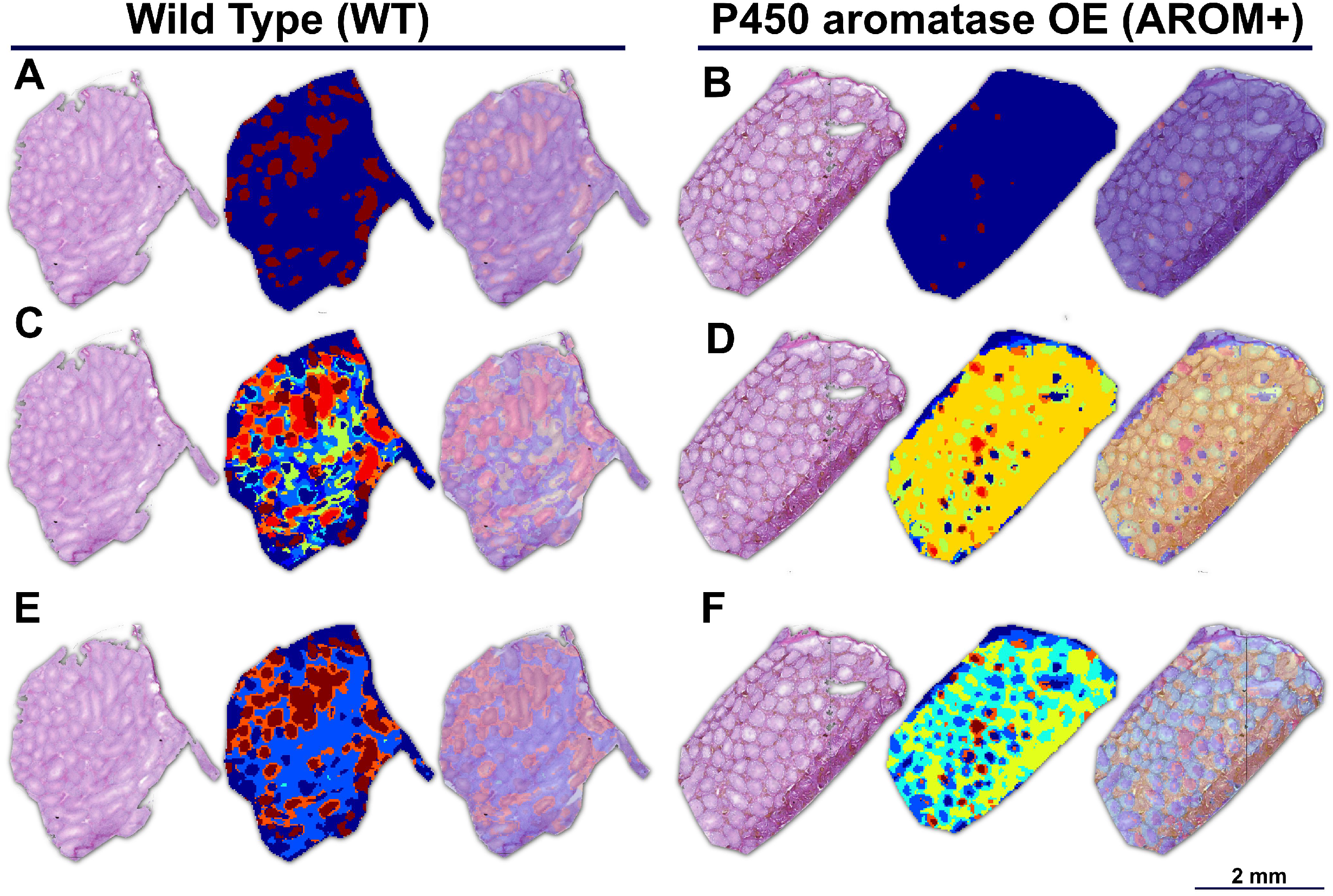
Proteomic map of WT and AROM+ testes. Initial unbiased hierarchical clustering of IMS peptide data from generated from whole tissue sections of **(A)** WT and **(B)** AROM+ testis (Dendrograms: Supplementary figure S1) at 25 μm spatial resolution. Further clustering results for the **(C)** WT and **(D)** AROM+ tissues (Dendrograms: Supplementary figure S2). In case of WT, 4 different sets of tubules are represented by the red, brown, green and deep blue clusters respectively. The orange cluster represents peritubular regions and the royal blue cluster characterizes the interstitial spaces. A further level of clustering of **(E)** WT and **(F)** AROM+ tissues (Dendrogram: Supplementary figure S3) shows an exclusive cluster (yellow-green) that is localized to the interstitial regions of AROM+ testis.

Detailed examination of the segmentation maps enabled us to further assign distinct spectral features to tubules (lumina and peritubular regions) or interstitial spaces (Fig 3C; red, brown, green, dark blue, orange and royal blue respectively). The tubular clusters along with their finer structures are less prominent in case of AROM+ (Fig 3D). Besides a general lack of a spectral diversity among the testicular tubuli in AROM+ mice, we identified an additional interstitial cluster in these animals compared to a WT control (Fig 3E and F yellow-green).

### Identification of proteins in situ

To get an idea about the parent proteins of the peptides representing different spectral clusters, we developed a bioinformatic pipeline that combines IMS and LC-MS/MS measurements using a maximum likelihood strategy (Figure 4).

**Figure 4:**
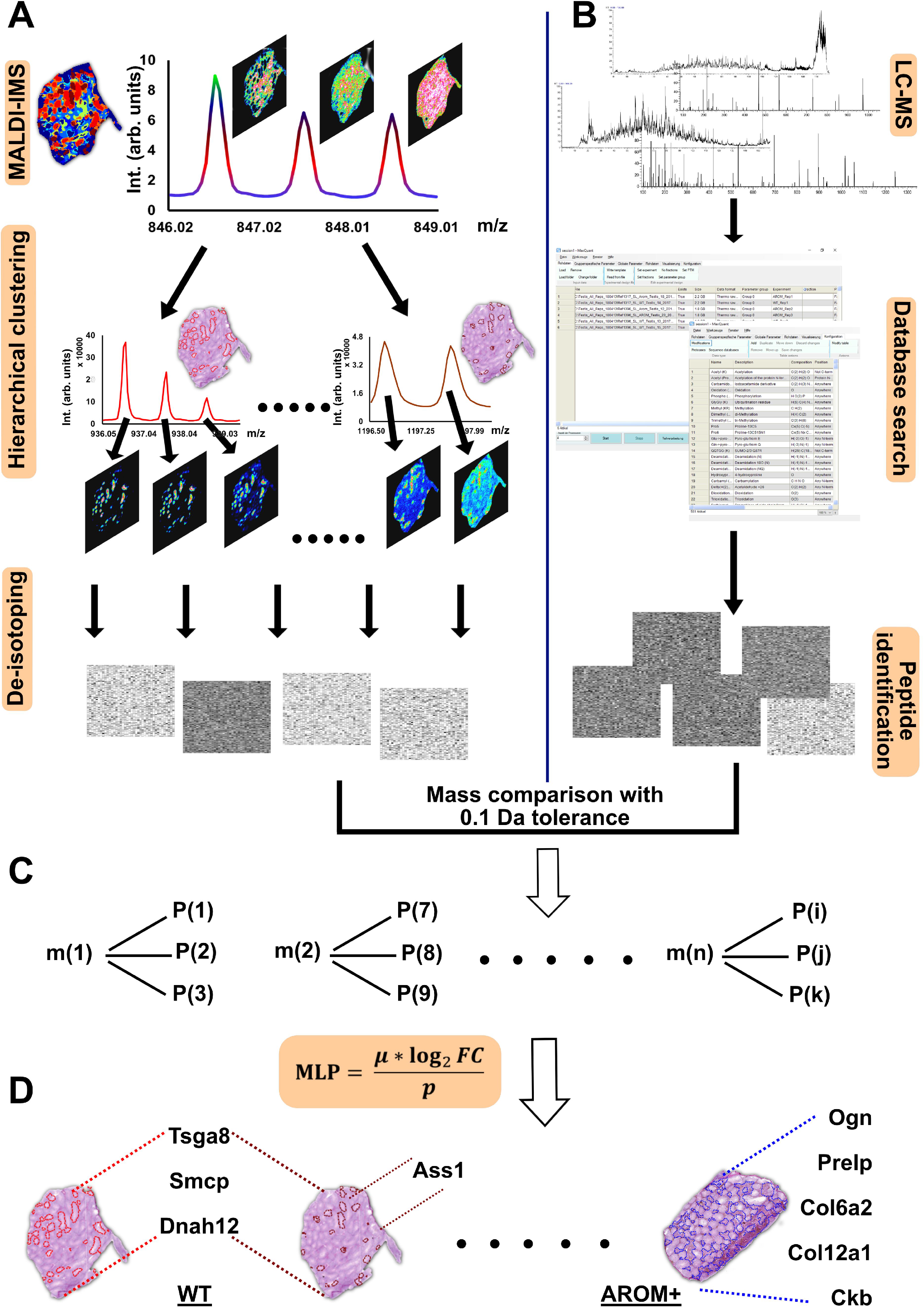
Protein identification *in situ* with maximum likelihood. **(A)** Imaging mass spectrometry (IMS) measurements lead to an overlap of isotope envelopes of different peptides in the same mass range (*Panel: MALDI-IMS*). This is indicated by different distribution patterns of the concerned m/z peaks. Therefore, to segregate the spectra, we have used unbiased hierarchical clustering (SCiLS Lab 2016b) of IMS data from the entire tissue (*Panel: Hierarchical clustering*) that led to efficient de-isotoping of the IMS spectra and create a mono-isotopic list (*Panel: De-isotoping*). **(B)** LC-MS/MS measurements were performed from tissue following the section used for IMS and peptides/proteins were identified following spectra analysis and database search (MaxQuant 1.6.0.16) **(C)** Upon mass comparison, each m/z value (m(1), m(2)…m(n)) from the IMS mono-isotopic list was annotated to multiple peptides identified in LC-MS/MS measurements (P(1), P(2)….P(k)) due to moderate mass accuracy (0.1 Da) of the current IMS set up. To clear the aforesaid ambiguity, we developed a scoring system (MLP scoring) that enabled to identify the parent proteins with maximum likelihood. **(D)** Distribution of identified proteins (proteins having highest MLP scores) in the WT and AROM+ testis. Proteins involved in late stage spermatogenesis (that were among the significantly altered proteins in LC-MS measurements also) were identified in the seminiferous tubules of WT (Supplementary file 7), whereas proteins involved in ECM regulation were found in the interstitia of AROM+ testis (Supplementary file 8).

The complexity of the tissue section used in MALDI-IMS results in multiple overlapping isotopic envelopes derived from the peptides present at a particular region within the tissue (Figure 4A, Panel: MALDI-IMS). This makes the deisotoping for any given peptide, which is required for all search algorithms extremely challenging and has been a major hurdle in MALDI-IMS based proteomics. To overcome this issue we took advantage of an unbiased hierarchical clustering of the entire imaging dataset (Figure 4A, Panel: Hierarchical clustering), which results in a segregation of the spectra based on their spatial distribution. As we expect all isotopes of a given petide to show the same spatial distribution, this strategy allowed us to separate overlapping isotopic envelopes with a very high accuracy. In mouse testis this enabled us to unambiguosly assign a regional distribtion to 369 unique deisotoped peptides from our MALDI-IMS spectra. To identify these peptides we compared this monoisotopic list with the peptides identified by LC-MS measurements of serial sections (Figure 4B). However, due to the moderate mass accuracy for peptide measurements (0.1 Da) this comparison often resulted in an ambiguous assignement of each m/z value to several peptides identified in the LC-MS analysis (Figure 4C). To overcome this ambiguity we devised a novel scoring method termed as ‘maximum likelihood peptide (MLP) score’. A particular m/z value from IMS was assigned to that identified peptide (from LC-MS analysis), which had the highest MLP score. Using this strategy we could identify masses discriminating the clusters from each other (Figure 4D).

The scoring could identify the spermatid associated the Axonemal dynein heavy chain 12 (Dnah12), sperm motility enabling Sperm mitochondrial-associated cysteine-rich protein (Smcp) and Testis-specific gene A8 protein (Tsga8) among others in the largest cluster (red) of tubules of WT mice (Figure 4D, Supplementary File 6). The next largest cluster (brown) shares the distribution pattern of the above mentioned proteins but in addition also contains peptide corresponding to arginosuccinate synthetase among the significantly enriched proteins in WT (Figure 4D, Supplementary File 7). As most of these proteins were shown to play a role in final stages of spermatogenesis and fertilization, we assume that the cluster corresponds to tubular stages IX-XI of spermatogenesis. Both clusters are barely detected in AROM+ mice (Figure 3D) supporting the hypothesis that aromatase over expression inteferes with late stages of spermatogenesis and fertilization.

The most prominent peptide defining the exclusive interstitial cluster of the AROM+ testis (Figure 4D) is derived from the extracellular matrix protein Mimecan (Ogn), a component of the keratan sulphate degradation pathway. Owing to the pixel based statistical specificity and sensitivity of the discriminative mass finding algorithm of SCiLS Lab program, it was not possible to get other distinctive masses for this cluster. However, MLP scoring of a manually curated mass list showing specific interstitial distribution in AROM+ led to the identification of proteins like different chains of collagen and prolargin involved in extracellular matrix assembly and regulation (Figure 4D, Supplementary File 8). The aforesaid proteins were found to be significantly enriched in AROM+ testis in the LC MS analysis (Supplementary file 3). Intersetingly, mimecan has been shown to be differentially regulated in men suffering from idiopathic germ cell aplasia (Alfano, Pederzoli et al., 2019).

### Loss of inter-tubular heterogeneity and tubule associated biological pathways in AROM+

A notable feature of the IMS measurements is a substantial inter-tubular heterogeneity (Fig 5A; red, brown, green and deep blue groups referred to as Tubuli A, B, C and D respectively from here on) in the WT tissue suggesting differential protein composition of the tubules at the same cross-section. Since different tubules in the same histological section of mouse testis represent different stages of spermatogenesis, it is likely that there are proteomic signatures correlating to these stages. In AROM+ tubules spermatogenesis is severely disrupted resulting in the absence of this heterogeneity and disruption of tubular stages (Fig 5A).

**Figure 5:**
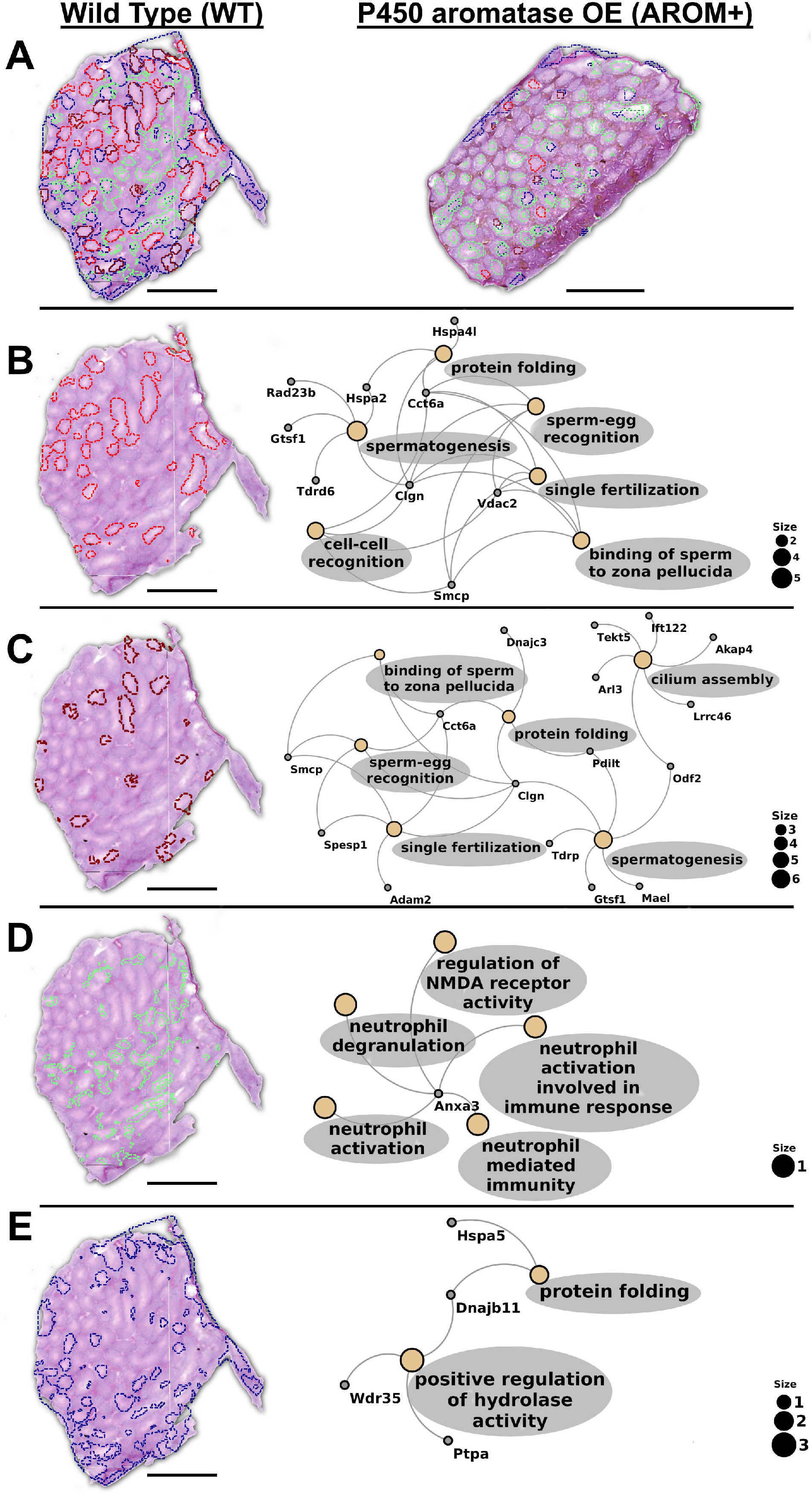
Inter-tubular heterogeneity and associated processes. **(A)** WT testis showing inter tubular heterogeneity (red, brown, green and deep blue) based on their protein composition. AROM+ testis (on the right side) does not exhibit this proteomic heterogeneity **(B)** GO term analysis of first set of tubules (red, Tubules A) based on the proteins identified there by MLP scoring. **(C)** GO term analysis of the second set of tubules (brown, Tubules B) based on the proteins identified there by MLP scoring. **(D)** GO term analysis of the third set of tubules (green, Tubules C) based on the proteins identified there by MLP scoring. **(E)** GO term analysis of fourth set of tubules (deep blue, Tubules D) based on the proteins identified there by MLP scoring. Scale bars – 1 mm GO term analysis was done based on Over-representation test that determines whether genes from belonging to a specific GO term are present more than that would be expected (over-represented) in a subset of the experimental data․. The yellow nodes in the network indicate the biological processes (named within the grey colored ovals). The size of these nodes in the GO term maps (scale represented by the numbered black solid circles) are representative of the number of genes involved in each of them. The grey nodes along with their labels show the genes that are involved in the aforesaid pathways.

A complete gene ontology (GO) analysis of the proteins identified in the MALDI-IMS data set by applying MLP scoring strategy shows that the two largest tubular clusters (A and B tubuli) are enriched for proteins involved in processes that are almost exclusively involved in late stages of spermatogenesis and fertilization (binding of sperm to zona pellucida, sperm-egg recognition, cell-cell recognition, single fertilization and cilium assembly and organization) (Figure 5B and 5C, Supplementary files 9a and 9b). Near absence of these clusters in AROM+ is in line with the data showing that tubules involved in late spermatogenesis and fertilization are severely disrupted in this model of male infertility. Tubuli C and D are observed to be present in relatively higher number in the AROM+ derived testis (Figure 3D). These tubuli are enriched for proteins involved in pro-inflammatory immune response and protein folding including ER associated unfolded protein response (Figure 5D and 5E, Supplementary Files 10a and 10b).

Therefore, it is observed that the healthy animals are characterized by sets of tubules with varied functions representing the complexity that is present and required for healthy spermatogenesis in testis. Impaired spermatogenesis, especially the late stages, in AROM+ leads to a severe reduction of this proteomic complexity, implying the necessity of region-specific molecular profiles in male gamete formation.

## DISCUSSION

Aromatase overexpressing (AROM+) mice are infertile and testicular changes closely resemble the ones observed in infertile men (Li et al., 2001). This testicular dysfunction is progressive in nature leading to severely impaired spermatogenesis in older mice (Li et al., 2006). To better characterize the molecular phenotype of these mice we performed an extensive proteomic analysis of the testes of 11-month-old AROM+ and WT mice combining bulk high-resolution shotgun proteomics with spatially resolving MALDI-IMS.

The comparative analysis of the bulk proteome of WT and AROM+ testes revealed several pathways misregulated in AROM+ mice. AROM+ for example show a significantly lower expression of proteins involved in androgen biosynthesis and metabolism of steroid hormones. Two of these (Cyp17a1 and Hsd17b3, Supplementary file 2) are Leydig cell markers, which have been shown to be down-regulated in the testis of AROM+ mice (Strauss et al., 2009). Such an impact of the aromatase induced increase in E2 levels may in fact be due to a severely changed mitochondrial metabolism, which is consistent with the loss of various mitochondrial pathways and associated proteins in AROM+ mice (Figure 2, Supplementary file 4). Energised and respiring mitochondria play an important regulatory role in LH-dependent steroidogenesis(Allen, Shankara et al., 2006), which is likely to be depleted in AROM+ mice despite normal levels of LH. This effect of increased aromatase levels on mitochondria is also consistent with the observation that mitochondria in Leydig cells of AROM+ are significantly enlarged(Strauss et al., 2009). In addition to the impaired steroidogenesis, we also found that testes of AROM+ mice express lower levels of estrogen sulfotransferase. This enzyme is abundantly expressed in Leydig cells and catalyses the formation of an estrogen sulfo-conjugate, thereby reducing the levels of active non-conjugated estrogen in the male testis(Qian, Sun et al., 2001). We therefore speculate that AROM+ mice have a higher E2/T ratio not only due to an increased E2 synthesis but also fail to endogenously synthesize androgens and are unable to compensate for high E2 levels by enzymatic conjugation, both of which are probably caused by a malfunction of Leydig cells.

Besides a number of proteins pointing towards an impairment of Leydig cell mitochondrial functions in AROM+ mice, we also detected lower levels of proteins directly involved in spermatogenesis such as the highly testis specific and GPI anchored protein Lypd4 (Bioinformatics) or late spermatogenesis factors like sperm mitochondrial-associated cysteine-rich protein, axonemal dynein heavy chain 12 and 17, phosphoglycerate kinase 2 and tubulin polyglutamylase TTLL5 (Supplementary file 1). These changes together with enrichment of male enhanced antigen1 in WT mice address the dysregulation of spermatid specific biological processes in development of the AROM+ phenotype (Ohinata, Sutou et al., 2002) resulting to lack of late stage spermatogenesis and spermatid specific processes. We also observe a role of hitherto unknown damping of compensatory mechanisms in development of the AROM+ phenotype.

In addition to proteins that are present at lower levels in AROM+ animals we also detect factors that are significantly upregulated upon aromatase overexpression. Most of these have been shown to be involved in the regulation of the extracellular matrix similar to what has been shown upon an upregulation of pro-inflammatory pathways (Adam, Urbanski et al., 2012, Li et al., 2006, Mayer, Adam et al., 2016, Walenta, Schmid et al., 2018).

Although being highly informative, such bulk proteomic analyses do not provide any information about the expression of a particular protein within the tissue. We therefore combined the bulk analyses with MALDI-IMS measurements of *in situ* digested peptides to gain insight into the molecular basis of tissue regions being altered upon higher E2/T ratio. Initial unbiased clustering of peptide masses show gross alteration of proteomic composition of seminiferous tubules pointing to a severe impairment of spermatogenesis in AROM+. Consistent with previous findings of interstitial fibrosis, we observe an interstitial space specific cluster in AROM+ absent in WT.

The combination of a comparative shotgun proteomics of the WT and AROM+ testes together with the MALDI-IMS measurements enabled us to identify the proteins to which these peptides defining particular *in situ* clusters belong. So far, this has been almost impossible by using either method separately as *in situ* generated peptides are low in number and can not be systematically fragmented to get sequence information and in turn, the bulk LC MS/MS experiments lack positional information. A combination of both has so far yielded either in very limited identification with considerable manual input or improved identification with resource intensive setups and limited applicability to different cell types (Alberts, Pottier et al., 2017, Huber, Khamehgir-Silz et al., 2018, Kriegsmann, Longuespée et al., 2017, Longuespée, Ly et al., 2019). We have therefore used peptide clustering together with a novel bioinformatic method (MLP scoring) to identify peptides, which can be applied at the MS1 level across all imaging platforms and tissues. This MLP based identification enabled us to associate relevant biological pathways with the most likely regions of the tissue. So are for example proteins associated with GO terms such as spermatogenesis, male gamete formation, etc. found within seminiferous tubules of WT whereas mimecan and collagen, two extra cellular matrix components were localized to the interstitial spaces of AROM+ testis.

In addition the scoring system also enabled us to localise proteins considerably enriched in WT. So does for example Calreticulin 3, a testis specific chaperone required for sperm fertility (Ikawa, Tokuhiro et al., 2011), localize to two largest set of seminiferous tubules taken together (Supplementary file 11) along with calnexin. Interestingly, asthenozoospermia causing sperm mitochondrial-associated cysteine-rich protein (Nayernia, Adham et al., 2002) is observed in the same tubuli and is not among the list of proteins enriched in AROM+ animals. We also observe a number of spermatogenesis related proteins like heat shock proteins, dipeptidase 3, outer dense fiber protein 2, tektin 4, etc. (Supplementary file 11) that are not significantly altered in the bulk analysis to be substantially reduced in AROM+ implying an additional and complementary coverage of molecular information that leads to better comprehension of the system.

Another interesting aspect of the combined analysis is the fact that we can use it to identify distinct sets of seminiferous tubules in the WT testis based on their proteomic composition. The set designated as A tubuli is defined by peptides derived from a series of proteins involved in fertilization related processes. These tubuli are almost absent in AROM+ suggesting an overall impairment of the late stages of spermatogenesis. Similar to the A tubuli, the B tubuli are characterized by factors associated with spermatogenesis, male gamete formation and associated metabolic processes and are disrupted in 11 month old infertile AROM+ mice. According to our MLP scoring approach, the set C and D tubuli are enriched in annexin, protein di-sulfide isomerase A6, endoplasmic reticulum chaperone BiP and its co-chaperone DnaJ homolog subfamily B member 11. These proteins are part of the immune response and ER associated unfolded protein response (UPR) in cells under stress when misfolded proteins accumulate in the ER (Chen & Brandizzi, 2013) respectively. This suggests that some seminiferous tubules of old (11 months age) mouse testis are already under cellular stress. Relatively higher presence of these cluster of tubules in AROM+ as compared to the type A and B tubuli suggests that AROM+ mice elicit an UPR during sperm maturation along with a triggering of pro-immune pathways. Although the cause for this increased UPR in AROM+ mice is unclear it might very well be that it induces a differentiation block and is therefore causative for the observed infertility.

In summary, we use a novel and unique bioinformatic strategy to identify peptides from IMS measurements based on a combination of bulk proteome analysis of healthy and diseased tissue with MALDI-IMS measurements. This method results in a reliable and high-throughput approach for molecular identification of peptides with maximum likelihood in MALDI-IMS. By applying this method to a model system of male infertility, we could achieve a deeper molecular understanding of the processes that cause this diseased phenotype. Along with corroborating previous findings, our study revealed new and unexplored proteins/pathways that are misregulated in male infertility. This technique opens up new avenues for further exploration of cellular processes as putative therapeutic targets not only for male infertility induced by hormone imbalance but also for multiple other diseases.

## Materials and methods

### Animals

Adult male mice were housed at Helmholtz Zentrum Munich (Neuherberg, Germany) under specific pathogen free conditions in GM500 cages, including individually ventilated caging systems (IVC System Green Line, Tecniplast, Buguggiate, Italy), which are operated with positive pressure. The mice were transferred to new cages with forceps in Laminar Flow Class II changing stations weekly; they were fed with an irradiated standard rodent high energy breeding diet (Altromin 1314, Altromin Spezialfutter GmbH & Co. KG, Lippe, Germany) and had access ad libitum to semi-demineralized filtered (0.2 mm) water. The light cycle was adjusted to a 12h/12h light/dark cycle; room temperature was regulated to 22 ± 1°C and relative humidity to 55 ± 5%. Husbandry conditions were adjusted to the experimental requirements in specified modules. Sentinels (outbred 8-week-old male SPF Swiss mice) were housed on a mixture (50:50) of new bedding material and a mixture of soiled bedding from all cages of this IVC rack and their health was monitored by on-site examination of certified laboratories according to the FELASA recommendations. Animal handling was in accordance with German and European guidelines for use of animals for research purposes. Experiments were further approved by the institutional animal care committee.

The AROM+ transgenic mouse was employed as a systemic model of inflammation-associated male infertility developing with increasing age. This mouse line expresses the human androgen-converting enzyme P450 aromatase under control of a universal promoter (ubiquitin C) in an FVB/N background (Li, Makela et al., 2003, Li et al., 2001, Li et al., 2006).

Testes were collected from ~320 days old AROM+ mice (n = 3) and age-matching wild-type littermates (n = 3).

### LC-MS/MS measurements

#### Sample preparation

Approximately 1 mg of tissue was incised from both WT and AROM+ mice testis. Initially the tissues were homogenized manually using Micro-homogenizers, PP (Carl Roth GmbH+Co. KG, Karlsruhe, Germany). Subsequently, proteins were extracted from the partially homogenized tissue employing the iST Sample Preparation Kit (PreOmics, Martinsried, Germany) using manufacturer’s protocol. Briefly; the tissues were lysed and subsequently sheared to get rid of DNA and other interfering molecules. The homogenous lysate was incubated with Trypsin at 37°C for two and a half hours. The ensuing peptides were subsequently desalted, purified and re-dissolved in 10 μl ‘Load’ solution after drying completely in a speed vac.

#### MS measurements

For reversed phase HPLC separation of peptides on a Ultimate 3000 nanoLC system (Thermo-Fisher-Scientific), 5 μl of the solution were loaded onto the analytical column (120 × 0.075mm, in house packed with ReprosilC18-AQ, 2.4 μm, Dr. Maisch GmbH), washed for 5 min at 300 nl/min with 3% ACN containing 0.1% FA and subsequently separated applying a linear gradient from 3% ACN to 40% ACN over 50 min. Eluting peptides were ionized in a nanoESI source and on line detected on a QExactive HF mass spectrometer (Thermo-Fisher Scientific). The mass spectrometer was operated in a TOP10 method in positive ionization mode, detecting eluting peptide ions in the m/z range from 375 to 1600 and performing MS/MS analysis of up to 10 precursor ions. Peptide ion masses were acquired at a resolution of 60000 (at 200 m/z). High-energy collision-induced dissociation (HCD) MS/MS spectra were acquired at a resolution of 15000 (at 200 m/z). All mass spectra were internally calibrated to lock masses from ambient siloxanes. Precursors were selected based on their intensity from all signals with a charge state from 2+ to 5+, isolated in a 2 m/z window and fragmented using a normalized collision energy of 27%. To prevent repeated fragmentation of the same peptide ion, dynamic exclusion was set to 20 s.

#### Data analysis

Protein identification was performed by MaxQuant 1.6.0.16 software package. Parent ion and fragment mass tolerances were 8 ppm and 0.7 Da respectively and allowance for 2 missed cleavages was made. Mouse canonical protein database from Uniprot (release June, 2018), filtered to retain only the reviewed entries was used for the searches. Regular MaxQuant conditions were the following: Peptide FDR, 0.01; Protein FDR, 0.01; Min. peptide Length, 5; Variable modifications, Oxidation (M); Acetyl (Protein N-term); Acetyl (K); Dimethyl (KR); Fixed modifications, Carbamidomethyl (C); Peptides for protein quantitation, razor and unique; Min. peptides, 2; Min. ratio count, 2. Proteins were validated on the basis of at least 1 unique peptide detected in the proteome of all the 3 replicates or in at least 2 of the 3 replicates.

#### Statistical analysis

In order to detect significant changes in protein levels between WT and AROM+, we performed statistical analysis on the acquired LC-MS/MS data. We chose LIMMA moderated t-test proposed earlier (Kammers, Cole et al., 2015) over standard t-test because it utilizes full data to shrink the observed sample variance towards a pooled estimate, resulting in far more stable and powerful inference compared to ordinary t-test.

Missing values in proteomics data is a common problem and are usually imputed before applying statistical methods. Here, we have used an imputation algorithm (Tyanova, Temu et al., 2016), which allows us to randomly draw values from a distribution meant to simulate values below the detection limit of the MS instrument. We have implemented the statistical analyses in R, MATLAB and used Cytoscape (Shannon, Markiel et al., 2003) for network graph generation.

#### Pathway analysis

In order to understand biological pathways affected in testis by the overexpression of aromatase, we analysed proteins that were either enriched or exclusively detected in WT or AROM+ for their involvement in those pathways. Enrichment analysis is a common practice to discover biological themes associated with the proteins of interest. Here, we used the R package ReactomePA (Yu & He, 2016) to discover biological pathways in which the enriched or exclusive proteins participate. The package uses hypergeometric distribution model to calculate p-values to determine whether any Reactome Pathways annotate a specified list of genes at a frequency greater than that would be expected by chance.

### Imaging mass spectrometry (IMS)

#### Sample preparation

Fresh frozen testis from WT and AROM+ animals (n = 3, in each case) were cryo-sectioned into 12 μm thick sections using CM1950 cryostat (Leica Microsystems; Wetzlar, Germany) and thaw-mounted on conductive indium-tin-oxide (ITO) coated glass slides (Bruker Daltonik GmbH; Bremen, Germany) pretreated with poly-lysine (1:1 in water with 0.1% NP-40). The slides were then stored at −80°C until further analysis.

Tissue sections were washed prior to trypsin and matrix spraying to remove lipids and salts, which act as interferents in MALDI measurements. Tissue sections were washed as previously described (Deutskens, Yang et al., 2011) with some modifications for measuring *in situ* generated peptides. Briefly; proteins were precipitated *in situ* by a sequential wash in 70% and 100% ethanol (for 30 s each) followed by a 90 s wash in Carnoy’s fluid (6:3:1 Ethanol:Chloroform:Acetic acid, v/v/v) to remove lipids and other interferants. Tissues were put back in 100% ethanol for 30 s to remove the excess chloroform. Further washing in dH_2_O containing 0.2% TFA was performed to remove salts and acidify. Excess acid was removed by washing the sections in 20 mM ammonium bicarbonate for 30 s. The excess water from the previous two steps was removed by washing with 100% ethanol for 30 s. Subsequently, the slides were dried in vacuum overnight at RT before Trypsin-spraying.

25 ng/μl Trypsin solution was prepared in 20 mM ammonium bicarbonate and sprayed on the tissue sections using the syringe spray system of TM Sprayer (HTX Imaging, HTX Technologies, LLC). The spraying parameters for trypsin were the following: Temperature - 30°C, No. of passes – 8, Pattern – Criss-Cross (CC), Flow rate – 0.03 ml/min, Nozzle velocity – 750, Track spacing – 2 mm, Gas pressure – 10 psi, Nozzle height – 40 mm. The sprayed sections were then incubated at 50°C for 2.5 hours in a humid chamber.

Subsequently, the slides were sprayed with 10 mg/ml HCCA in 70% ACN and 1% TFA in the TM sprayer using the following spray parameters: Temperature – 75°C, No. of passes – 4, Pattern – Horizontal (HH), Flow rate – 0.12 ml/min, Nozzle velocity – 1200, Track spacing – 3 mm, Gas pressure – 10 psi, Gas flow rate: 2, Nozzle height – 40 mm.

#### IMS measurements

Imaging experiments were performed in a rapifleX MALDI Tissuetyper MALDI-TOF/TOF mass spectrometer (Bruker Daltonik GmbH; Bremen, Germany) equipped with a SmartBeam 3G laser. Tissues were measured in a positive reflector mode with 25 μm spatial resolution and using a mass range of 600-3200 Da. Each measurement was pre-calibrated externally using a commercial peptide calibrant mixture (Bruker Daltonik GmbH; Bremen, Germany) spotted on the same target as the tissues at multiple positions. Tissues from all the animals were measured in the rapifleX using 1.25 GS/s sample rate and a pulsed ion extraction of 160 ns.

After the IMS measurements, HCCA matrix was removed by washing the slides in 70% ethanol and H&E counterstaining was performed. High-resolution images of the stained sections were obtained from Mirax Scan system (Carl Zeiss MicroImaging, Munich, Germany) and co-registered with MALDI-IMS data for histological correlation.

#### Data Analysis

The acquired data was exported to SCiLS Lab version 2016b (SCiLS, Bremen, Germany, www.scils.de) (Trede, Schiffler et al., 2012) software for further analysis. Initially, a segmentation pipeline was executed on the acquired IMS data from both WT and AROM+ tissues for unbiased hierarchical clustering with a minimal interval width of ± 0.1 Da. The resulting clusters were examined for their correlation with histological features of the tissues manually. Subsequently, clusters representing relevant regions were selected for further analysis.

Peptide masses defined as distinguishing the clusters that are discussed in the manuscript were found using‘Find discriminative m/z values’ tool of SCiLS Lab. The tool uses the Receiver Operating Characteristic (ROC) measure to discriminate between all m/z values from a particular cluster. The resulting mass list in each case (with a threshold of at least 0.7) was exported as an excel file and further processed through in-house developed program (as described in the following section).

#### Analysis and identification of peptide masses

The goal was to identify proteins based on IMS peptides at the MS1 level with maximum likelihood. To achieve this, we performed multistage data analysis. The acquired IMS data was deisotoped taking advantage of co-localisation of peptides in different clusters. Deisotoping was performed on these co-localised peptides using standard tolerances (for m/z (± 0.15) and for intensity (50%)). Each mass in the deisotoped IMS mass list was then matched with the LC-MS peptide masses (after adding the mass of a proton due to singly charged ions in MALDI) within a tolerance τ. The choice of τ depends on the accuracy of IMS measurements and in this case τ = ± 0.1 was appropriate. LIMMA statistics (as explained before) was applied on the LC-MS peptide data (‘peptides’ output file from MaxQuant analysis) and the peptides for the two groups WT and AROM+ were classified on the basis of fold change (log_2_FC), defined as

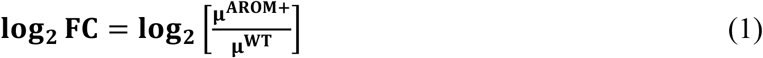

Where, μ^AROM+^ and μ^WT^ imply mean intensity of a particular peptide across the AROM+ and WT replicates respectively. All peptides that have log_2_FC > 0 were classified as AROM+ and those that have log_2_FC < 0 were classified as WT peptides. Subsequently, IMS masses were compared with either WT and AROM+ data depending on relative abundance of peptides in the tissues.

However, for one IMS peptide mass, the τ-search method yielded multiple LC-MS peptides of similar masses (± 0.1 Da.) belonging to different proteins. Therefore, in order to determine the maximum likelihood of the peptide belonging to a particular protein, we devised a scoring system based on the following assumptions/conditions:

a. Peptides of relatively higher abundance are preferably detected in MALDI-TOF-IMS since there are no pre- or post-ionisation separation techniques involved. Therefore, it is more likely for a peptide detected in IMS to belong to an abundant protein than to its possible relatively less abundant counterpart.
b. The search space for a peptide belonging to a cluster detected in IMS measurements would be restricted to either WT or AROM+ LC-MS data depending on the occurrence of the cluster in the respective tissues.

The aforesaid scoring system (MLP) is then defined as follows:

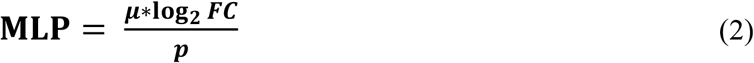

Where, μ is the mean intensity of a particular peptide across 3 animals, log_2_ *FC* is the fold change of that peptide for either WT or AROM+ and *p* imply LIMMA moderated p-value of the same peptide. All these parameters were extracted from the LC-MS data of WT and AROM+ tissues. Therefore, higher the MLP score of an IMS peptide mass for a particular protein, more likely it is to belong to that protein. For all the IMS peptides, we then select the proteins with the highest MLP score, which are subsequently used to systematically investigate biological processes and pathways to which they are associated.

We performed GO enrichment analysis by the means another R package clusterProfiler(Yu, Wang et al., 2012) to identify the enriched GO categories associated with proteins. This package uses hypergeometric distribution to calculate p-values to determine whether any GO terms annotate a specified list of genes at a frequency greater than that would be expected by chance.

#### Gene Ontology (GO) Enrichment Analysis

In order to find out the biological processes and networks represented by the identified proteins in situ, we performed GO based enrichment analysis using Over-representation test (Boyle, Weng et al., 2004) as implemented in clusterProfiler (Yu et al., 2012). In this, we assessed whether any gene ontology-biological process (GO-BP) term annotates our list of proteins at a frequency greater than that would be expected by chance.

#### Software availability

The codes for LIMMA statistics with data visualization, the τ-Search method and MLP scoring modules will be available at the following GitHub link - https://github.com/wasimaftab/IMS_Shotgun

## ABBREVIATIONS

MALDI-IMS: Matrix assisted laser desorption/ionisation imaging mass spectrometry
LC-MS: Liquid chromatography mass spectrometry
WT: Wild type
AROM+: Mice overexpressing human P450 aromatase
E2/T: Estrogen/Testosterone ratio
ECM: Extracellular matrix
ER: Endoplasmic reticulum
GO: Gene ontology
UPR: Unfolded protein response

## Acknowledgements

The authors thank Dr. Juan Antonio Aguilar-Pimentel from Helmholtz Zentrum Munich for providing the mice. Thanks are also due to the Group of Prof Dr. Doris Mayr, Pathologisches Institut der LMU München for their enormous help with cryo-sectioning of tissues for IMS measurements. We thank Dr. Ignasi Forne of the Protein analysis unit (ZfP) at BMC, LMU for his help with the LC-MS measurements. Thanks to Ms. Irene Vetter, Molecular Biology department, BMC, LMU for her excellent assistance in sample preparation for IMS measurements and post-measurement staining of the testis cross-sections. This work was supported in part by DFG grants MA1080/23-1; −2, 27-1and 29-1 (to AM), the DAAD/Academy of Finland (DAAD project 57347353; to AM, MP). This work was supported in part by DFG grants MA1080/23-1; −2, 27-1 and 29-1 (to AM), CRG1309 and CRG1123 (to AI) and the DAAD/Academy of Finland (DAAD project 57347353; to AM, MP).

## Author contributions

SL, AI and AM conceptualised the project and designed the experiments. SL and LW prepared the samples. SL performed the experiments. WA wrote and developed the codes used in the bioinformatic pipeline. SL, WA and AI analysed the data. SL and WA prepared the figures. SL, AI, AM, LS, MP, WA and LW discussed the findings and wrote the manuscript.

